# Early life stress induces social behavioral deficits and peripheral biomarker alterations in adolescence that perpetuate intergenerationally

**DOI:** 10.1101/2025.11.27.690841

**Authors:** Josephine C. McGowan, Jed L. Katzenstein, Natalie S. Chace, Camila Wagner-Sanchez, Temi Okotore, Tolu Ogunfowora, Tallie Z. Baram, Oluwarotimi O. Folorunso, William A. Carlezon

**Affiliations:** Department of Psychiatry, Harvard Medical School, Boston, MA, USA, 02115; Basic Neuroscience Division, Department of Psychiatry, Harvard Medical School, McLean Hospital, Belmont, MA, USA; Current address: Putnam Associates, New York, NY, USA; Jordan High School, Fulshear, TX, USA; Department of Neuroscience, University of Michigan, MI, USA; Center for Biomedical Engineering, Brown University, Providence, RI, USA; Department of Pediatrics, University of California, Irvine, CA, USA; Department of Neurology, University of California, Irvine, CA, USA; Department of Anatomy & Neurobiology, University of California, Irvine, CA, USA

**Author notes:** Corresponding Author: William A. Carlezon, Jr., Ph.D. Address: McLean Hospital of Harvard Medical School 115 Mill Street Mailman Research Center Belmont, MA 02478 Co-Corresponding Author: Oluwarotimi O. Folorunso, Ph.D. Address: McLean Hospital of Harvard Medical School 115 Mill Street Mailman Research Center Belmont, MA 02478.

**Keywords:** early life stress, maternal behavior, ultrasonic vocalizations, social interaction, stress markers, intergenerational

## Abstract

Early life stress (ELS) increases the likelihood of developing chronic health problems including mental illness. In humans, deficits in social behavior often emerge during adolescence and correlate with later-life psychiatric diagnoses. We examined in mice the effects of a limited bedding and nesting (LBN) model of ELS on pup ultrasonic vocalizations (USVs) during postnatal days 3-9 (P3-P9). Then on P30, an adolescent timepoint, we assessed social behavior as well as thymus involution and adrenal hypertrophy, both biomarkers of stress. We found reductions in USVs as early as P3, resembling low levels normally seen later in development. There were few changes in SI, with deficits observed only after restraint stress in female LBN mice. Surprisingly, thymus weights were augmented and adrenal glands were smaller in LBN adolescent mice, opposite to alterations typically observed after chronic adult stress. LBN also produced signs of precocious puberty in both sexes, especially in cohorts of LBN-exposed offspring bred to create second-generation LBN offspring that subsequently underwent LBN, indicating perpetuation across generations. Together, these data suggest that stress in early life has distinct and diverse effects, including accelerating several processes, and that some of these effects persist intergenerationally.

## INTRODUCTION

Accumulating evidence indicates that early life stress (ELS) has profound and persistent effects on health. For example, individuals who experienced chronic stressors during early development such as neglect, abuse, and poverty are more likely to experience increased inflammation during adulthood^1^, which can lead to myriad health issues across the lifespan, including increased susceptibility to mental health problems (e.g., anxiety, depression)^1–3^, cardiovascular and metabolic diseases^4–6^, cancer^7^, and neurodegenerative diseases^8^. ELS has been shown to have lifelong impacts on memory and executive functioning and is associated with increased prevalence of depression, anxiety, and post-traumatic stress disorder (PTSD)^9^. Abnormal social interaction is often an early sign of risk of developing psychiatric disorders^10^. Additionally, social interaction difficulties are a hallmark of many neuropsychiatric conditions such as autism spectrum disorder (ASD), attention-deficit/hyperactivity disorder (ADHD), and social anxiety disorder^11,12^. Often, these conditions emerge during adolescence, which has been shown in both rodents and humans^3,13^, representing a particularly vulnerable period as well as a critical timepoint for intervention before symptoms develop further into adulthood. Yet, whether there are specific behavioral or biological indications of susceptibility to social behavioral deficits prior to adolescence remains unknown. Characterizing when and how the effects of ELS manifest behaviorally and physiologically before they fully develop can lead to predictive indicators of psychiatric disease, ultimately driving initiation of targeted treatments at critical timepoints in the clinic.

Given the role of social behavior in psychiatric disease, we sought to determine the impact of ELS on putative measures of social communication^14,15^ and interaction^10^ during rearing and adolescence and determine their relationship to peripheral indices of stress. Chronic exposure to stress causes characteristic changes in peripheral organs and blood-borne biomarkers. For example, chronic stress causes hypertrophy (increased size) of the adrenal glands, which are a key node in the hypothalamic-pituitary-adrenal stress (HPA) axis, and involution (shrinkage) of the thymus, which plays a central role in regulation of immune system function and inflammation^16,17^. While these processes can be uncoupled, suggesting overlapping but distinct mechanisms^18^, they are accompanied by changes in blood markers that can serve as easily-accessible proxies that enable *in vivo* analysis of stress effects over time. Blood levels of corticosterone (CORT) are associated with adrenal gland size and function, and circulating proinflammatory cytokines (e.g., interleukin-6 [IL-6]) or TRECs (T-cell receptor excision circles) can provide insight on inflammatory processes and thymic function. TRECs are a novel biomarker—circular DNA biproducts of *de novo* T-cell production in the thymus—that correlate with stress history in both mice and humans and are strongly associated with potentially destructive processes such as accelerated epigenetic aging^18^. Because they are easily accessible and scalable across species, use of these biomarkers in mice to reveal susceptibility to stress and reflect the long-lasting impact of ELS can help to enhance alignment of research performed in humans and animal models.

Evidence shows that the consequences of stress can also be transmitted via intergenerational and transgenerational epigenetic inheritance. One well-documented example is the Hunger Winter of the Netherlands in 1944, which caused alterations in gene function, DNA methylation, and elevated stress responses in subsequent generations that persisted throughout life^19,20^. More recent evidence found intergenerational transmission of elevated cortisol and PTSD risk in offspring of mothers pregnant during the 9/11 attacks^21^. While there is compelling evidence in *C. elegans*, *Drosophila*, and plants that inheritance of stress effects occurs through small RNAs and histone modifications^22–25^ which cause lasting epigenetic changes in offspring, mechanistic studies of these effects in mammals have not yet been comprehensively evaluated^26–29^.

To reproduce ELS in mice in the present study, we used the limited bedding and nesting (LBN) model developed by the Baram laboratory^30–34^, which recapitulates scarcity and poverty during the early postnatal period, leading to fragmented, maternal behavior in the dam as well as lasting brain, behavioral, and endocrine system dysregulation in the offspring^35,36^. Pups were reared in either control cages rearing or LBN cages during postnatal days (P) 2–9. Second-generation LBN mice were produced by pairing LBN-exposed mice, then exposing their offspring to LBN for subsequent experiments. We then analyzed social behavior during the rearing period by recording ultrasonic vocalizations (USVs), a putative measure of social communication. During neonatal development, mouse pups typically communicate with their mothers using vocalizations in the ultrasonic range. First coined as “whistles of loneliness”^37^, pup USVs serve as an early communicative behavior of the mother-pup dyad: USVs trigger maternal care and facilitate communication between mother and offspring^38–40^. USVs thus provide insight into the emotional state of mouse pups as well as reflect modifications in the arousal states of the mother^38,41^. Mice then underwent a social interaction (SI) test on P30, considered an adolescent timepoint in mice^42–45^. Our data indicate that ELS acutely alters social behavior but does not itself produce a behavioral phenotype into adolescence.

However, we uncovered robust peripheral changes, specifically in stress-relevant organs and pubertal timing, that indicate ELS has a lasting impact on the normal process of maturational development into adolescence as well as future stress susceptibility, which can transmit intergenerationally.

## MATERIALS AND METHODS

### Mice

Male and female C57BL/6J mice were obtained from Jackson Laboratory at 8 weeks of age (Jackson Laboratories, Bar Harbor, ME) and bred in-house in a controlled vivarium on a 12 h:12 h (0600h:1800h) light cycle, with *ad libitum* access to food and water. Experiments were approved by the McLean Hospital Institutional Animal Care Use Committee (IACUC) and carried out according to National Institutes of Health (NIH) guidelines for experimental animals. All efforts were made to minimize animal suffering and the number of animals used.

### Limited Bedding and Nesting (LBN)

The LBN paradigm was performed as previously described^31,46^. Briefly, dams were assigned to control (CTRL) or LBN conditions, which was kept consistent for up to 5 litters from the dams. Litters with fewer than 4 pups were excluded and litters larger than 8 were culled based on sex to maintain a balance between males and females. CTRL dams were placed into standard cages containing a normal amount of Alpha Chip (Northeastern Products Corp., Warrensburg, NY) bedding (∼1 inch thick distributed evenly across the cage floor) and one cotton nestlet. In LBN cages, the bedding was reduced to scarcely cover the cage bottom to absorb urine (ammonia), a fine gauge aluminum mesh was placed ∼2.5cm above the cage floor, and half of a cotton nestlet was provided (50% reduction). From P2-9, cages were left undisturbed, except during USV recording sessions. Maternal videos were recorded each day from 0600-0800 and 1800-2000. On P10, all mice were ear-clipped for identification and placed back to standard cages with normal bedding and nesting. For tracking experiments, mice were labeled for identification using a microtattoo system and green permanent ink (Fine Science Tools, Foster City, CA) at P2 on the top of their paws before changing the cage to their CTRL or LBN rearing conditions. On P10, all cages reverted to standard cages with normal bedding and nesting.

To generate second generation LBN pups, male and female pups that had undergone LBN (but no other experimental condition) from different litters were randomly chosen and paired together. Their litters were then also subjected to LBN according to the same timeline as described above. Maternal behavior for a subset of litters across groups was quantified at P6 during 1h of the active period (lights-off) and 1h of the inactive period (lights-on) using the behavioral observation research interactive software (BORIS)^47^. Maternal events recorded included: active nursing (AN), carrying pups (C), eating (E), licking and grooming pups (LG), low nursing (LN), moving on the nest (M), nest building (NB), moving off the nest (O), self-grooming (SG), side nursing (SN), and tail biting / chasing (TB), a stress-related rodent behavior that has been previously reported^48^. Time that pups were off the nest was also scored.

### Puberty tracking

One cohort of pups was used to monitor pubertal status, as previously reported in rats^49^. Briefly, mice were assessed for vaginal opening in females and preputial separation in males every day after weaning from P21-P30 or whenever they reached pubertal completion. Visual inspection was used, during which mice were picked up individually by an experimenter and inspected for 15-30 seconds per day. For males, pubertal initiation was determined by initial preputial separation, defined as any separation of the prepuce from the glans penis. Full puberty was defined as the ability to fully retract the prepuce. For females, pubertal initiation was defined as the appearance of a vaginal pinhole, which typically appears ∼2 days before full vaginal opening.

### Statistical analyses

All analyses were performed using Prism 10.5.0 (GraphPad Software, La Jolla, CA). The Shapiro-Wilk test was used to assess the normality of the data, followed by *t*-tests, Mann-Whitney U tests, or either 1-way or 2-way analyses of variance (ANOVAs) with a Dunnett’s or Holm–Sidak tests for *post-hoc* analyses. Pearson correlation tests were used for all correlation analyses. Statistical outliers were identified using the Grubbs test and excluded from analysis. The threshold for statistical significance was set at *p* < 0.05. All statistical analyses are reported in **Supplemental Table 1**.

Detailed methods are provided in the **Supplement.**

## RESULTS

### LBN induces fragmented maternal behavior that perpetuates across generations

We first sought to validate that the LBN model of ELS reliably induces fragmented maternal behavior, and that the behavior transmits across generations. We conducted home cage maternal observations in a randomly selected subset of litters, quantifying the number of behavioral transitions during postnatal day 6 (P6) specifically during 1 h of both the active (lights-off) and inactive (lights-on) periods. Total duration of behaviors as well as the number of transitions to each behavior was quantified using BORIS^47^. Four mice per condition were chosen at random from dams that produced litters for the present experiments. Both LBN (F(2,15)=12.11, *p*=0.0029) and perpetuated LBN (LBN 2G) dams (F(2,15)=12.11, *p*=0.0007) exhibited a higher number of behavioral transitions and a higher probability of transitioning to other behaviors (**Fig. 1A-D; Suppl. Fig. 1-3**). Specifically, the duration (LBN: F(2,15)=6.482, *p*=0.0412; LBN 2G: F(2,15)=6.482, *p*=0.0063) and number (LBN: F(2,15)=4.697, *p*=0.0486; LBN 2G: F(2,15)=4.697, *p*=0.0239) of tail biting behaviors were increased in LBN and LBN 2G dams (**Fig. 1E-F; Suppl. Video 1**). The average duration of licking and grooming was significantly decreased in LBN and LBN 2G dams (LBN: F(2,15)=15.3, *p*=0.0003; LBN 2G: F(2,15)=15.3, *p*=0.0007) (**Fig. 1G**). Additionally, types of self-directed behaviors, adverse pup-directed behaviors, and off-the-nest behaviors were altered differentially during the inactive and active phases of the experiment, with the largest effects in the combined data (**Fig. 1H-J; Suppl. Fig. 1-3; Suppl. Videos 2-3**). There were no differences in litter size across groups (F(2,49)=0.7143, *p*=0.4946) (**Suppl. Figure 4**). Overall, we recapitulated past work demonstrating that the LBN model induces fragmented, maternal behaviors as described before^31,50^ and have established that many of the adverse maternal behaviors perpetuate across generations in the LBN 2G dams.

**Figure 1.**
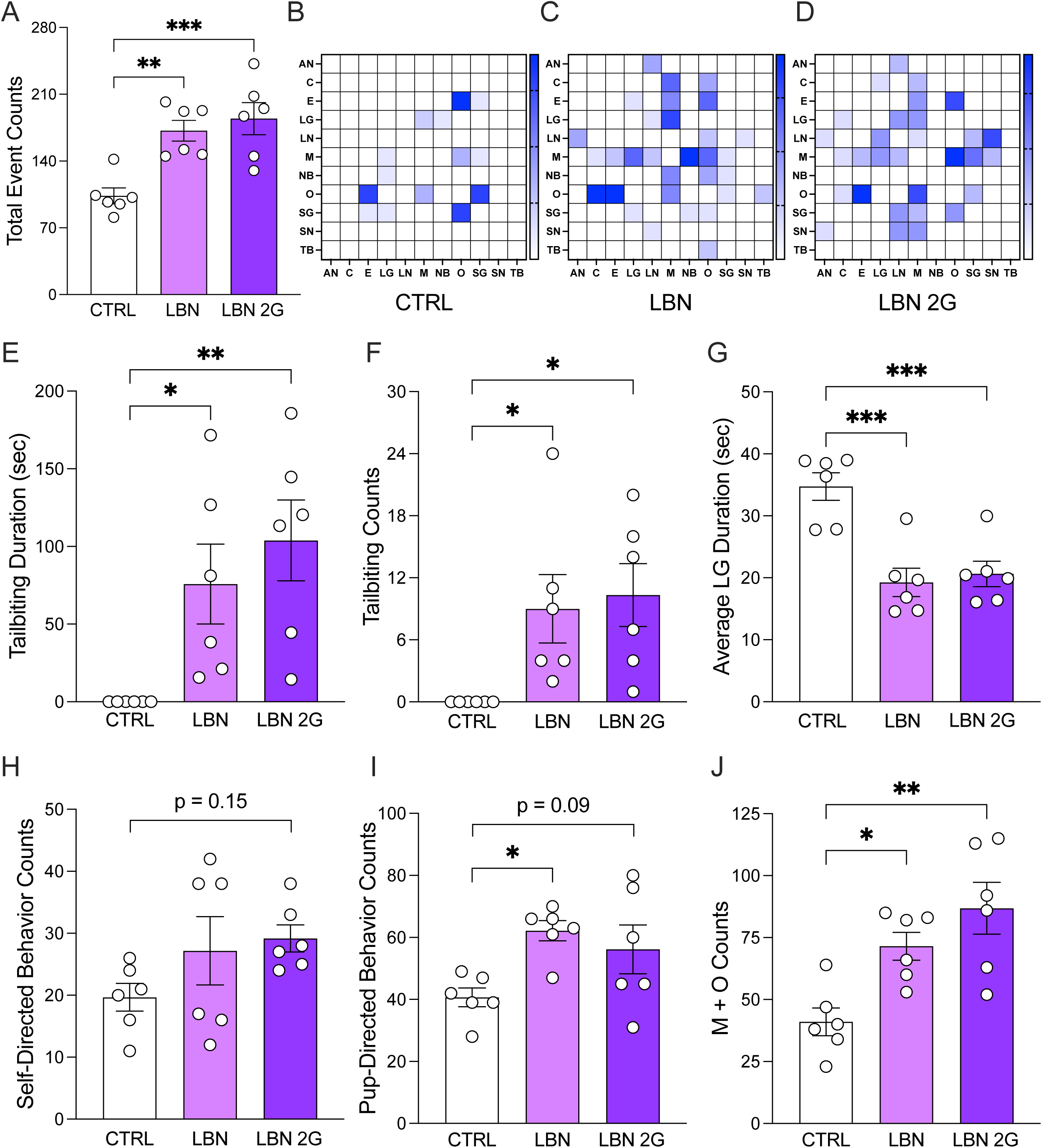
The limited bedding and nesting model of early life stress induces fragmented maternal behavior which transmits across generations. (A) Total behavioral event counts in both the active and inactive phases of postnatal day 6 (P6) is significantly higher in both LBN and LBN 2G dams. (B-D) Representative heatmaps depicting transition matrices from CTRL, LBN, and LBN 2G dams. Each box represents a color-mapped scale of the probability of moving from one behavior to another, with deeper blues representing higher probabilities of behavioral transitions, and thereby higher fragmentation of behavior. (E-F) LBN and LBN 2G dams engage in significantly more tail biting, a behavior that was not observed in CTRL conditions. (G) LBN and LBN 2G dams demonstrate increased frequency of licking and grooming pups. (H) There is a trend towards an increase in self-directed behaviors (self-grooming and eating) in LBN 2G dams as compared to CTRL dams. (I) LBN dams exhibit a greater number of pup-directed behaviors (nursing, nestbuilding, licking and grooming pups, and carrying pups) as compared to CTRL dams, with a trending increase in LBN 2G dams. (J) LBN and LBN 2G dams demonstrate significantly higher counts of moving on and off the nest as compared to the CTRL dams. (n =6 dams per group). Error bars represent ± SEM. * *p* < 0.05, ** *p* < 0.01, *** *p* < 0.001. CTRL, control; LBN, Limited Bedding and Nesting; LBN2G, LBN 2nd generation; M, Moving on Nest; O, Off the Nest, AN, Active Nursing; C, Carrying Pups; E, Eating; LG, Licking and Grooming Pups; LN, Low Nursing; NB, Nestbuilding; SG, Self-Grooming; SN, Side Nursing; TB, Tail biting; sec, seconds.

### LBN alters pup-dam social communication during rearing but not during adolescence

We first aimed to understand the effect of LBN on social attachment between the pup and dam during the rearing period. Mice first underwent either CTRL or LBN rearing from P2-P9. At P3, P6, and P9, the dam was separated from the pups for 10 minutes and USVs of pups were recorded one at a time with an ultrasonic microphone in a Styrofoam sound chamber for 3 minutes (**Fig. 2A**). LBN (F(2,49)=21.31, *p*<0.0001) and LBN 2G (F(2,49)=21.31, *p*<0.0001) (**Fig. 2B**) mice weighed significantly less than their control counterparts by P9 (**Fig. 2C,D**). LBN (F(2,46)=30.35, *p*<0.0001) and LBN 2G (F(2,46)=30.35, *p*<0.0001) mice exhibited decreased call number as early as P3, with effects persisting until P6 in LBN 2G mice (*p*=0.0278) (**Fig. 2E,F**). LBN mice exhibited longer call lengths on P6 as compared to CTRL (*p*=0.0001) or LBN 2G (*p*=0.0011) mice (**Fig. 2G,H**). The bandwidth of calls increased in both LBN (F(2,48)=6.855, *p*=0.0024) and LBN 2G LBN (F(2,48)=6.855, *p*=0.0304) mice on P9 (**Fig. 2I,J**). Spectrograms obtained from USV recordings revealed higher bandwidth and length of calls in LBN-reared animals (**Suppl. Fig. 5**). These data indicate that the LBN paradigm leads to lower numbers of USVs, but the calls are longer and occur over a more diverse range of frequencies through the rearing period, and this effect perpetuates intergenerationally.

**Figure 2.**
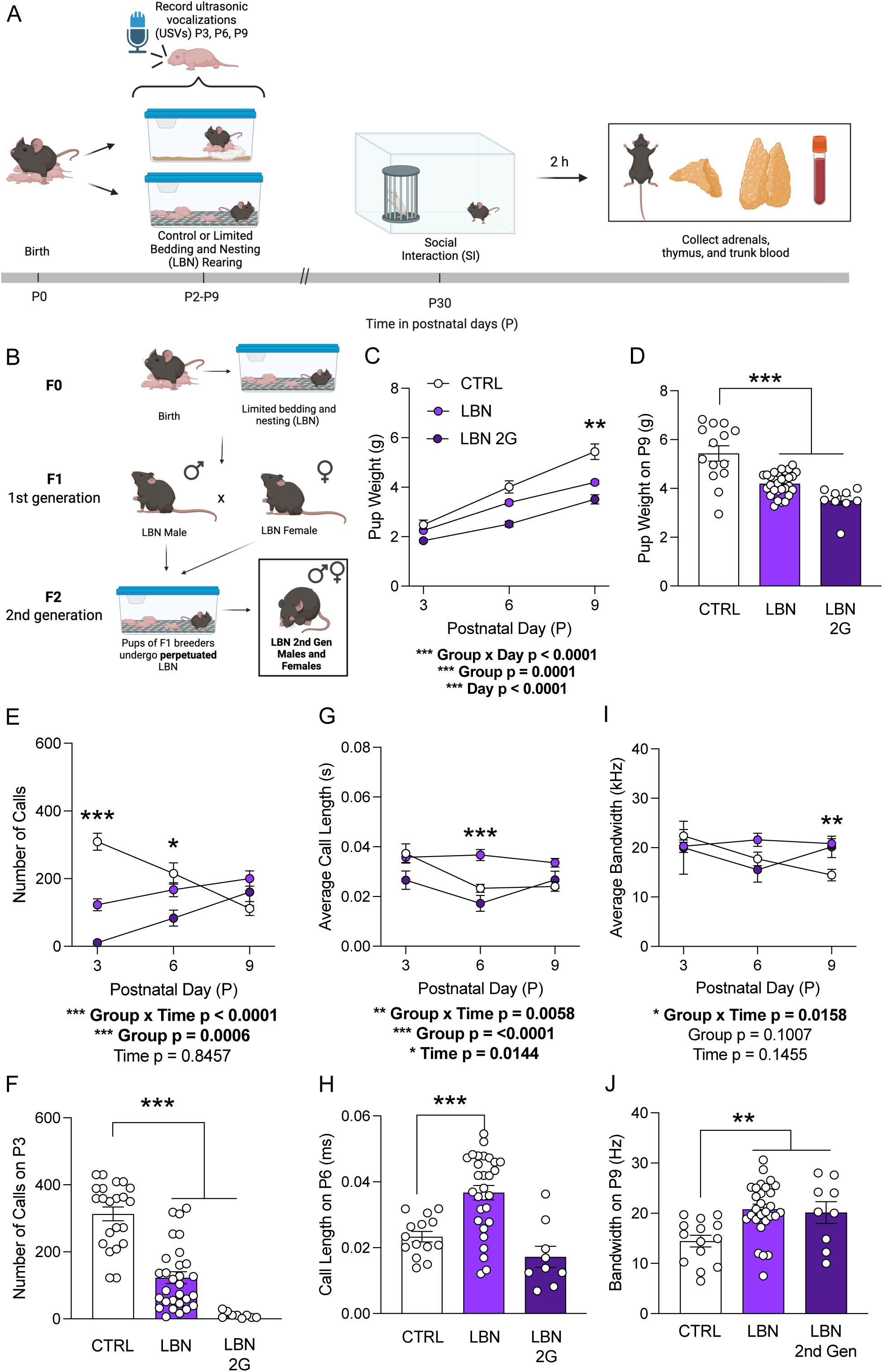
Early life stress induces social behavioral alterations via pup-dam communication during the early postnatal period and intergenerationally. (**A**) Experimental design. (**B**) Breeding paradigm for LBN 2G mice. (**C**) LBN and LBN 2G pups weigh less than CTRL pups across the rearing period. (**D**) On P9, LBN and LBN 2G mice weigh significantly less than CTRL mice. (**E-F**) The number of pup USV calls is significantly lower in the LBN and LBN 2G groups on P3 and P6. (**G-H**) Call length is significantly increased in LBN pups on P6. (**I-J**) The average bandwidth of pup calls is significantly higher on P9 in both LBN and LBN 2G pups as compared to CTRL mice. (n = 9-29 mice per group). Error bars represent ± SEM. * *p* < 0.05; ** *p* < 0.01; *** *p* <0.001. CTRL, control; LBN, Limited Bedding and Nesting; LBN 2G, LBN 2nd generation; P, postnatal day; USV, ultrasonic vocalization.

We next assessed social behavior in these mice at P30, which captures the period of pubertal onset and roughly corresponds to the teen years in humans^43^. There were no robust differences in social interaction in male mice (LBN: F(2,31)=1.71, *p*=0.857; LBN 2G: F(2,31)=1.71, *p*=0.1423) (**Fig. 3A**), but there were fewer bouts to the stimulus mouse only in LBN 2G male mice (F(2,31)=2.98, *p*=0.0468) (**Fig. 3B**). There were no differences in social interaction in female mice (LBN: F(2,25)=1.312, *p*=0.3322; LBN 2G: F(2,25)=1.312, *p*=0.3249) (**Fig. 3C**), though there was a trend towards a decrease in bouts to the stimulus mouse only in LBN 2G female mice (F(2,24)=1.845, *p*=0.1877) (**Fig. 3D**). The time spent with the cup, time spent with the mouse, total distance traveled, and average velocity did not significantly differ across groups in all mice (**Suppl. Fig. 6**). Overall, these data reveal that while LBN induces alters social behavior in the early postnatal period, there are no robust deficits in social interaction that persist to adolescence.

**Figure 3.**
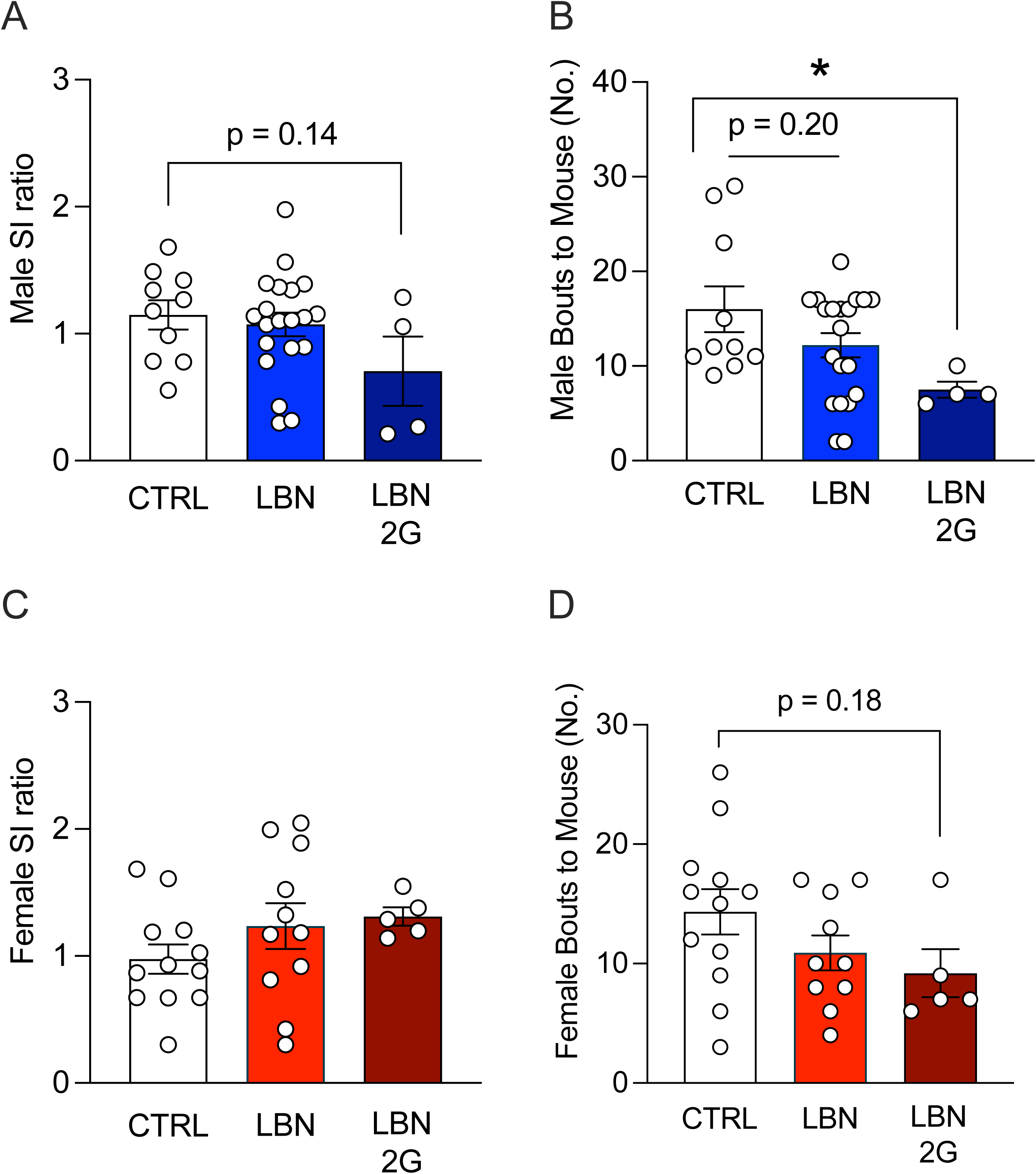
Early life stress does not impact social interaction during adolescence, but there is an intergenerational deficit in males. (**A**) Male SI ratio does not differ across groups at P30, though there is a trend towards a lower SI ratio in LBN 2G males. (**B**) The number of bouts to the stimulus mouse is significantly decreased in LBN 2G males, with a trend towards a decrease in LBN male mice as compared to CTRL male mice. (**C**) Female SI ratio does not differ across groups at P30. (**D**) The number of bouts to the stimulus mouse trends lower in LBN 2G female mice as compared to CTRL female mice. (n = 4-20 mice per group). Error bars represent ± SEM. * *p* < 0.05. SI, social interaction; CTRL, control; LBN, Limited Bedding and Nesting; LBN 2G, LBN 2nd generation; P, postnatal day; No., number.

### Pup-dam communication during rearing does not predict social behavior in adolescence after LBN

Because there were no differences in social interaction behavior during adolescence, we next examined whether social communication during rearing predicts sociability in adolescence. We utilized a paw tattoo system on P2 to track the behavior of the same mouse from P2 to P30 (**Fig. 4A**). This analysis also allowed us to also assess sex differences during the rearing period. During rearing, the overall number of USV calls were significantly lower at P3 in both males (F(2,24)=3.706, *p*=0.024) and females (F(2,42)=8.619, *p*=0.0003), replicating findings from **Fig. 2** (**Suppl. Fig. 7A**). These effects persisted in female LBN mice to P6 (F(2,41)=4.189, *p*=0.0466) and in LBN 2G mice to P9 (F(2,42)=10.38, *p*=0.0013) (**Suppl. Fig. 7B,C**). The overall number of USV calls throughout the rearing period decreased in LBN mice (Male: F(2,24)=10.34, *p*=0.0006; Female: F(2,42)=19.19, *p*<0.0001) but was not altered in LBN 2G mice (Male: F(2,24)=10.34, *p*=0.445; Female: F(2,42)=19.19, *p*=0.3131) (**Suppl. Fig. 7D**). We next replicated our finding that LBN does not induce SI deficits in adolescence (F(5,58)=1.38, *p*=0.2453) (**Fig. 4B**). The time spent with the cup, time spent with the stimulus (social partner) mouse, bouts to the mouse, total distance traveled, and average velocity also all did not differ across groups in adolescence (**Suppl. Fig. 8**).

**Figure 4.**
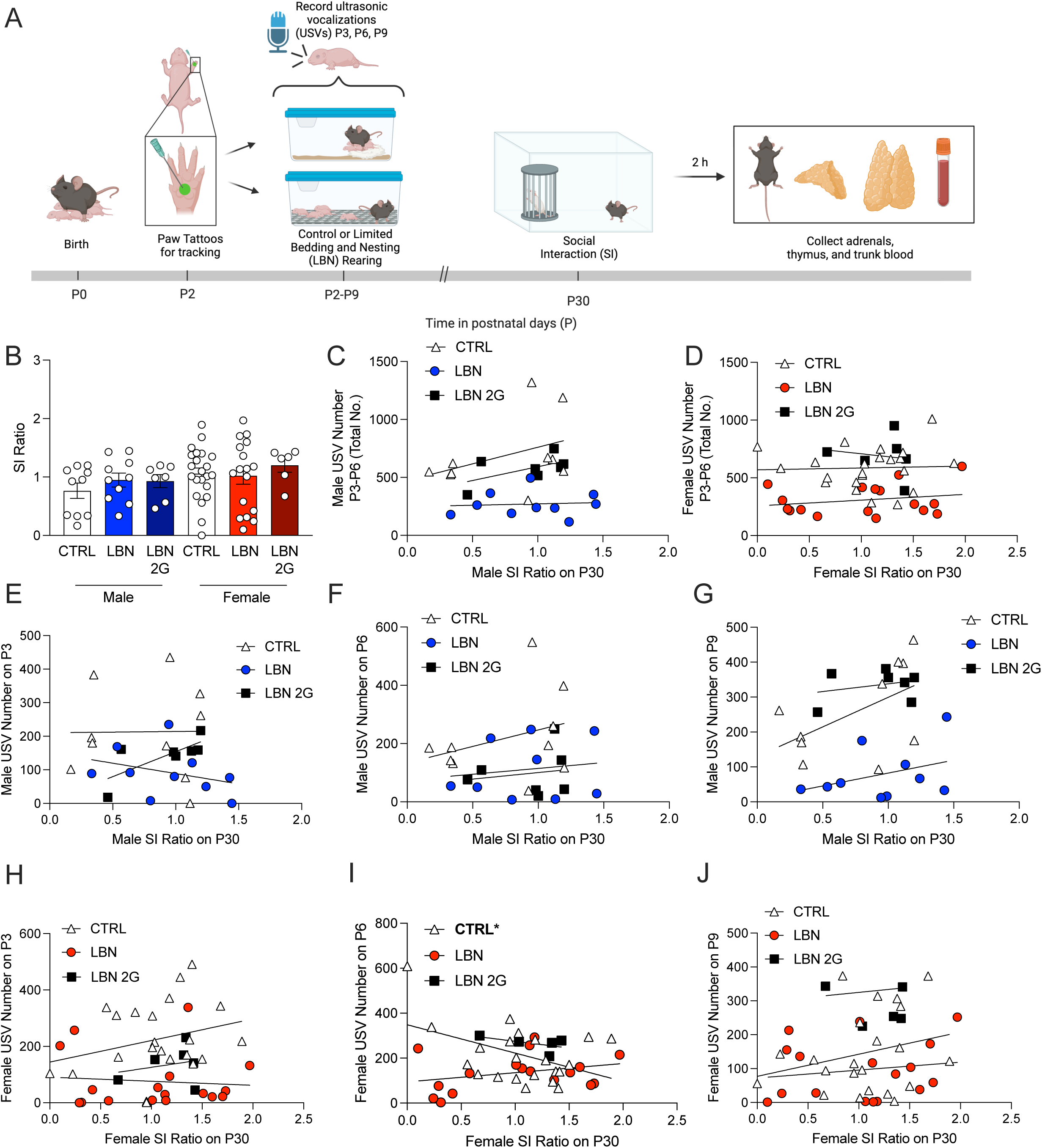
Ultrasonic vocalizations during rearing do not predict social interaction in adolescence after early life stress or intergenerationally. (**A**) Experimental timeline. (**B**) There are no effects on SI ratio across groups. (**C-D**) There is no significant correlation between total USV number (P3, P6, and P9 combined) and SI ratio on P30 in either males or females. (**E-G**) There are also no significant correlations between USV number on P3, P6, or P9 and SI ratio on P30 in males. (**H-J**) There is a significant negative correlation between USV number on P6 in CTRL females and SI ratio on P30, but no other correlations reach significance. (n = 6-22 mice per group). Error bars represent ± SEM. * *p* < 0.05. USVs, ultrasonic vocalizations; SI, social interaction; CTRL, control; LBN, Limited Bedding and Nesting; LBN 2G, LBN 2nd generation; P, postnatal day; No., number.

Correlating USV number to SI ratio, we found that the total number of USVs throughout the rearing period did not significantly predict SI behavior in either males (CTRL M: F(1,8)=1.333, *p*=0.2816; LBN M: F(1,8)=0.0526, *p*=0.8244); LBN 2G M: F(1,5)=0.3362, *p*=0.1724) or females (CTRL F: F(1,20)=0.03324, *p*=0.8572; LBN F: F(1,15)=0.7957, *p*=0.3865); LBN 2G F: F(1,4)=0.1042, *p*=0.7631) across groups (**Fig. 4C,D**). There were also no significant correlations revealed in the relationship between the number of USV calls on each day and SI ratio, though there was a positive correlation uncovered at P6 only in CTRL females (F(1,20)=4.986, *p*=0.0371) (**Fig. 4E-J**). Overall, these data reveal that USV number during these early timepoints in rearing does not significantly predict SI behavior in adolescence.

### LBN induces alterations in a sex-dependent manner which perpetuates across generations

While there were no behavioral differences revealed in the SI in adolescence, we next determined whether peripheral alterations occurred as a result of the LBN paradigm and perpetuated across generations, focusing on stress-relevant organs (thymus, adrenals) as well as blood-borne markers of stress and inflammation (corticosterone [CORT] and interleukin-6 [IL-6]).

In male mice, LBN caused decreased body weight in adolescence (F(2,68)=16.24, *p*=0.0039), which perpetuated intergenerationally (F(2,68)=16.24, *p*<0.0001) (**Fig. 5A**). In male mice, thymus weight (adjusted for body weight: mg/kg) significantly increased in both LBN (F(2,66)=9.679, *p*=0.0030) and LBN 2G (F(2,66)=9.679, *p*=0.0002) (**Fig. 5B**).

**Figure 5.**
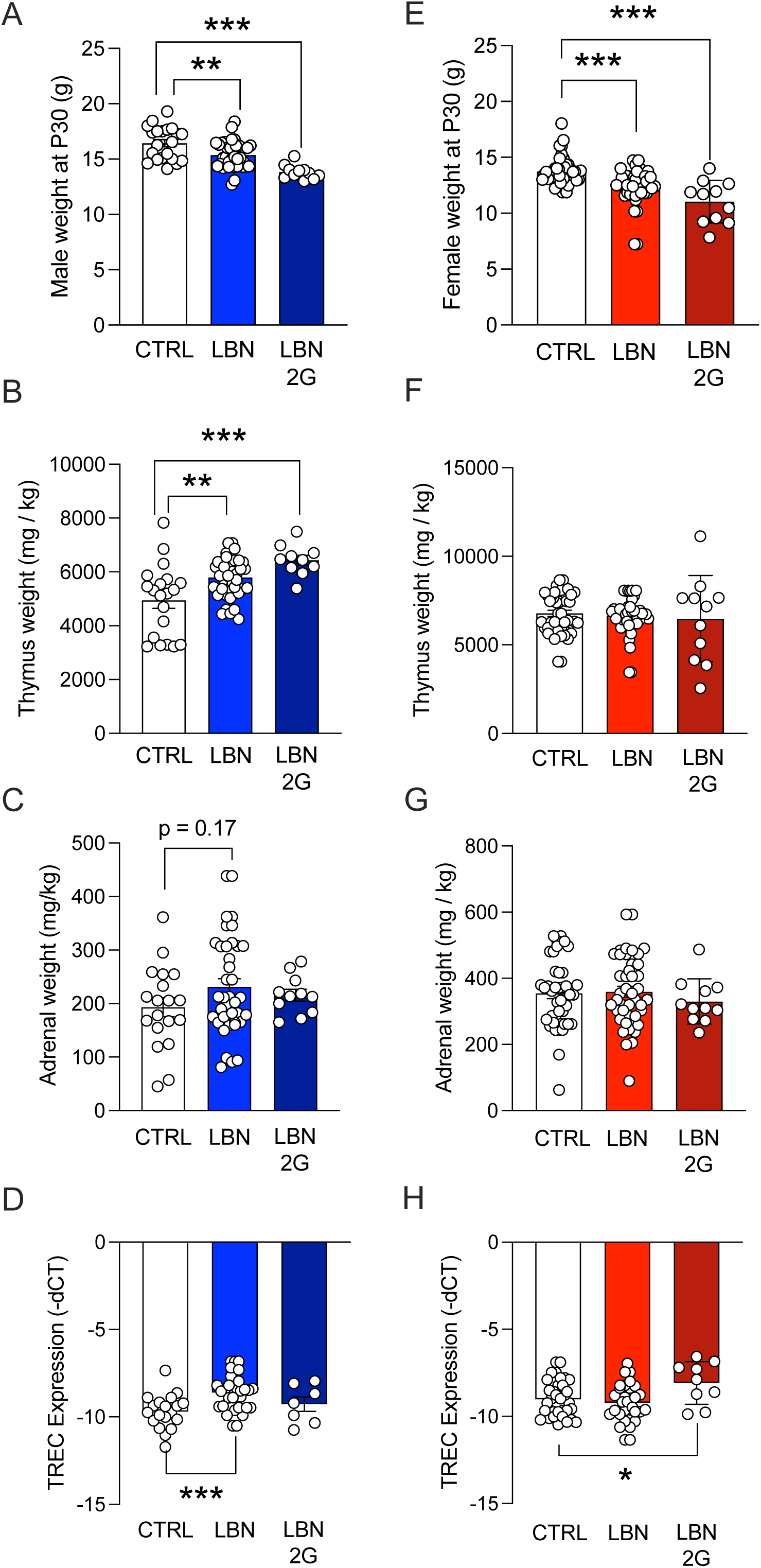
Stress-relevant organs and body weights are significantly altered in adolescence after early life stress and intergenerationally. (**A**) LBN and LBN 2G males weigh significantly less than CTRL males at P30. (**B**) LBN and LBN 2G male thymus weights are significantly higher than those of CTRL males. (**C**) LBN male adrenal weights trend higher than those of CTRL males. (**D**) TREC expression is significantly higher in LBN male mice as compared to CTRL males, while there are no differences in TREC expression in LBN 2G male mice. (**E**) LBN and LBN 2G females weigh significantly less than CTRL females at P30. (**F**) There is no difference across female groups in thymus weights. (**G**) There is no difference across female groups in adrenal weights. (**H**) TREC expression is significantly higher in LBN 2G female mice as compared to CTRL females. Note that TREC data are depicted as -dCT to clarify that higher dCT integer values indicate lower levels of TRECs. (n = 7-44 mice per group). Error bars represent ± SEM. * *p* < 0.05; ** *p <* 0.01; \*\*\**p* < 0.001. CTRL, control; LBN, Limited Bedding and Nesting; LBN 2G, LBN 2nd generation; mg/kg, milligrams per kilogram; -dCT, negative change of cycle counts between TREC gene expression and control gene expression.

Similarly, adrenal weights (mg/kg) trended towards an increase in LBN male mice (F(2,66)=1.423, *p*=0.1703) (**Fig. 5C**). However, non-body-weight-normalized thymus or adrenal weights did not differ across groups in males (**Suppl. Fig. 9A,B**). Consistent with increased thymic size, blood levels of T-cell receptor excision circles (TRECs), a measure of thymic output, were higher in LBN mice (F(2,57)=7.394, *p*=0.0007), although no differences were seen in LBN 2G mice (F(2,57)=7.394, *p*=0.5485) (**Fig. 5D**).

In female mice, LBN caused decreased body weight in adolescence, which perpetuated intergenerationally (F(2,96)=18.94, *p*<0.0001) (**Fig. 5E**). Thymus weight (mg/kg) was not altered across groups (LBN: F(2,96)=0.2879, *p*=0.8395; LBN 2G: F(2,96)=0.2879, *p*=0.7293) (**Fig. 5F**), though non-body-weight-normalized thymus weight in LBN 2G female mice decreased (F(2,65)=5.874, *p*=0.0036) (**Suppl. Fig. 9C**). Similarly, adrenal weights (mg/kg) did not differ across groups (LBN: F(2,95)=0.1348, *p*=0.9948; LBN 2G: F(2,95)=0.1348, *p*=0.8398) (**Fig. 5G**) or when analyzed as non-normalized over body weight (**Suppl. Fig. 9D**). Interestingly, blood levels of TRECs were higher in LBN 2G female mice (F(2,71)=3.833, *p*=0.0478) (**Fig. 5H**), suggesting an uncoupling between thymic size and weight that may reflect changes in the rate of TREC degradation and turnover^51^.

Male LBN mice exhibited increases in CORT in the blood (*p=*0.0145), while LBN 2G male mice exhibited a blunting of the CORT response (*p=*0.0.8766) (**Suppl. Fig. 10A**). Female mice did not exhibit alterations in CORT expression across groups (LBN: *p*=0.9965; LBN 2G: *p*=0.9494). Blood plasma levels of IL-6 trended towards an increase in LBN-reared mice (F(2,26)=16.05, *p*<0.0001), with the most significant increase being in LBN 2G female mice (*p*<0.0001) (**Suppl. Fig. 10B**). Overall, these peripheral data reveal that there are robust, sex-dependent alterations in body weight, thymus weight and output, and blood-borne biomarkers of stress and inflammation after LBN.

### A brief stressor in adolescence reveals social behavior deficits only in LBN female mice

Considering the peripheral alterations observed after LBN during adolescence, we hypothesized that this underlying biology may indicate altered vulnerability to stress at this timepoint. To test this hypothesis, we exposed CTRL and LBN-reared mice to a brief 10-minute restraint stress 2 hours before SI at P30 to induce a stress response (**Fig. 6A**). Interestingly, only female mice administered both LBN and restraint stress exhibited an SI deficit (*p*=0.0408), as compared to control mice that underwent restraint, revealing a sex-specific stress vulnerability-like effect (**Fig. 6B,C**) reflected by reductions in the total time spent with the stimulus mouse (F(2,34)=4.95, *p*=0.0228) (**Suppl. Fig. 11D**). There were no differences in time spent with the cup, time spent with the stimulus mouse, bouts to the mouse, total distance traveled, or average velocity during the SI across all other groups (**Suppl. Fig. 11**). Both LBN (F(2,22)=10.18, *p*=0.08) and LBN 2G male mice (F(2,22)=10.18, *p*=0.0005) exhibited a decrease in weight, though for this experiment there were no differences in the weight of female mice across groups (LBN: F(2,33)=0.1576, *p*=0.9296; LBN 2G: F(2,33)=0.1576, *p*=0.8383) (**Fig. 6C**). The thymus weight over body weight was increased in LBN 2G males (F(2,24)= 3.392, *p*=0.0345) but decreased in LBN 2G females (F(2,33)=6.561, *p*=0.004) (**Fig. 6D**), findings which were reflected when analyzing non-normalized weights (**Suppl. Fig. 12A,C**). Adrenal weights did not differ across male groups but significantly decreased in LBN 2G female mice (F(2,33)=10.08, *p*=0.0002) as compared to CTRL female mice (**Fig. 6E**), findings which were reflected when analyzing non-normalized weights (**Suppl. Fig. 12B,D**). Representative images of a thymus (male) and adrenal glands (female) for each experimental group are shown in **Suppl. Fig. 13**. There is an increase in TREC expression in the blood in LBN 2G female mice (F(5,53)=2.633, *p*=0.0249) (**Fig. 6F**). Restraint stress did not increase CORT in LBN mice as expected but rather blunted it in both LBN 2G male (F(5,38)=9.493, *p*=0.0015) and female (F(5,38)=9.493, *p*=0.0002) mice (**Fig. 6G**). Overall, these data suggest that LBN females exhibit a vulnerability to adolescent stress for social behaviors, and that there are marked blunting of stress-induced CORT intergenerationally in both sexes during adolescence.

**Figure 6.**
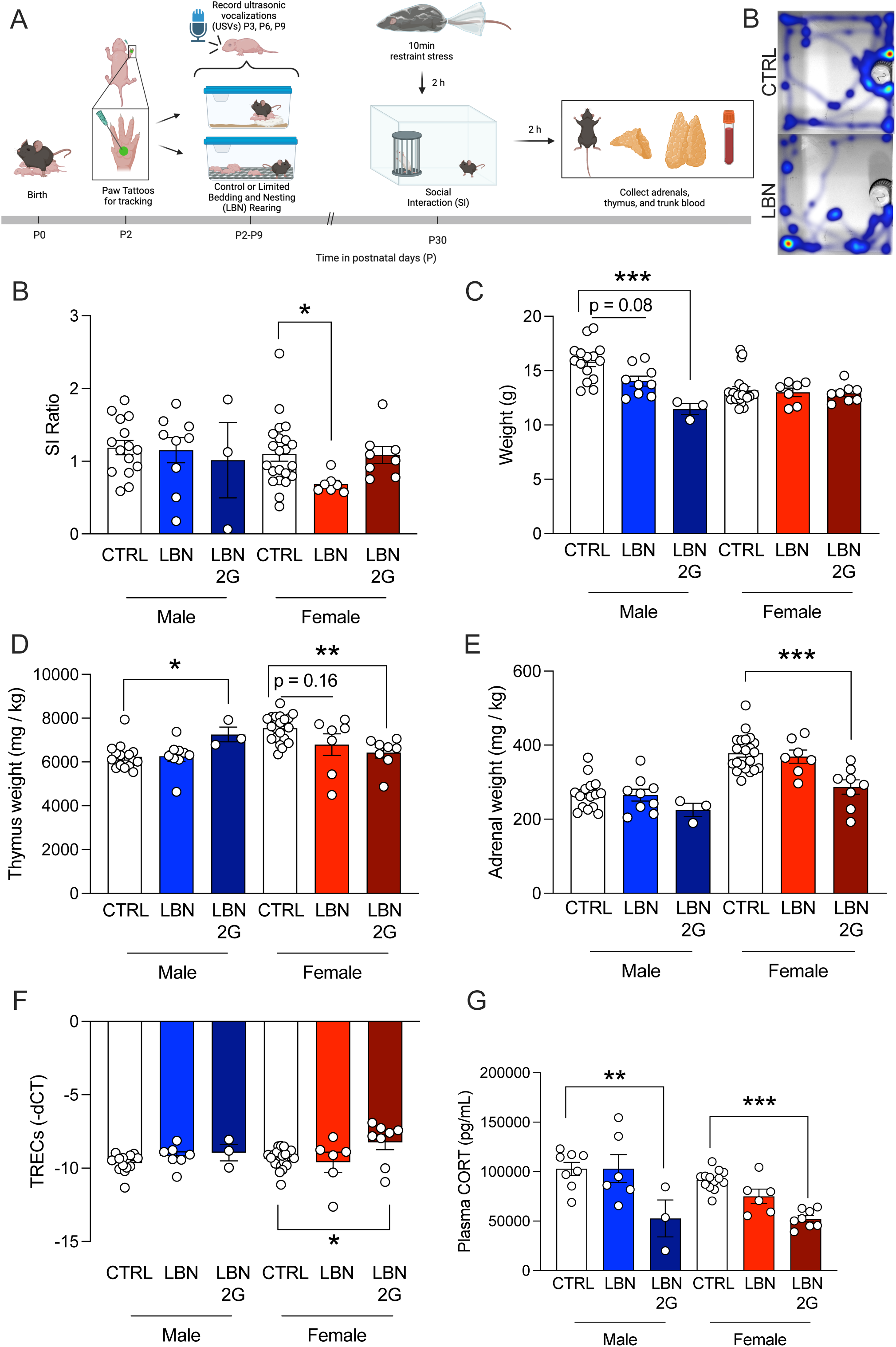
A brief restraint stress during adolescence after early life stress significantly impacts social interaction in female mice. (A**) Experimental timeline.** (**B**) SI ratio is significantly decreased only in LBN females as compared to CTRL females. (**C**) Male LBN 2G mice weigh significantly less than CTRL males, with a trend towards decreased weights in LBN males. (**D**) LBN 2G males have a larger thymus weight as compared to CTRL males, while LBN 2G females have a decreased thymus weight as compared to CTRL females, with LBN females trending towards decreased weights. (**E**) LBN 2G female mice have significantly smaller adrenal size than CTRL females. (**F**) There is no difference in TREC expression in male groups. TREC expression is significantly higher in LBN 2G female mice as compared to CTRL females. Note that TREC data are depicted as -dCT to clarify that higher dCT integer values indicate lower levels of TRECs. (**G**) Plasma CORT is significantly lower in LBN 2G males and females as compared to CTRL mice. (n = 7-44 mice per group). Error bars represent ± SEM. * *p* < 0.05; ** *p <* 0.01; \*\*\**p* < 0.001. USVs, ultrasonic vocalizations; CTRL, control; LBN, Limited Bedding and Nesting; LBN 2G, LBN 2nd generation; g, grams; mg/kg, milligrams per kilogram; -dCT, negative change of cycle counts between TREC gene expression and control gene expression; pg/mL, picograms per milliliter.

### LBN causes earlier initiation of puberty but not completion in males and females

Considered together, the data above reveal several phenotypes that may indicate either early or late maturation. The thymus normally peaks in weight in adolescence around P40 in mice^16,52,53^, indicating a potential accelerated maturation in LBN mice, while body weight differences across experiments indicate a potential delay in maturation. To determine whether LBN accelerated or delayed maturation, we next assessed pubertal onset, an indicator of early adolescence and an indication of biological aging^54^. Separate groups of mice were evaluated from P23 to P30+ for signs of pubertal initiation and completion. Mice were observed once every day for preputial separation (males) and vaginal opening (females). We found that both male (LBN M: F(5,42)=31.27, *p*=0.0095; LBN 2G M: F(5,42)=31.27, *p*<0.0001) and female (LBN F: F(5,42)=31.27, *p*=0.0031; LBN 2G F: F(5,42)=31.27, *p*<0.0001) LBN-reared mice initiated puberty earlier (**Fig. 7A**) but completed puberty at the approximately the same rate (LBN M: F(5,42)=7.343, *p*=0.571; LBN 2G M: F(5,42)=7.343, *p*=0.9999; LBN F: F(5,42)=7.343, *p*=0.7405; LBN 2G F: F(5,42)=7.343, *p*=0.1984) (**Fig. 7B**), indicating earlier overall pubertal timing in both sexes. These results suggest that LBN induces alterations in maturational processes, specifically earlier sexual maturation, during adolescence that perpetuates across generations.

**Figure 7.**
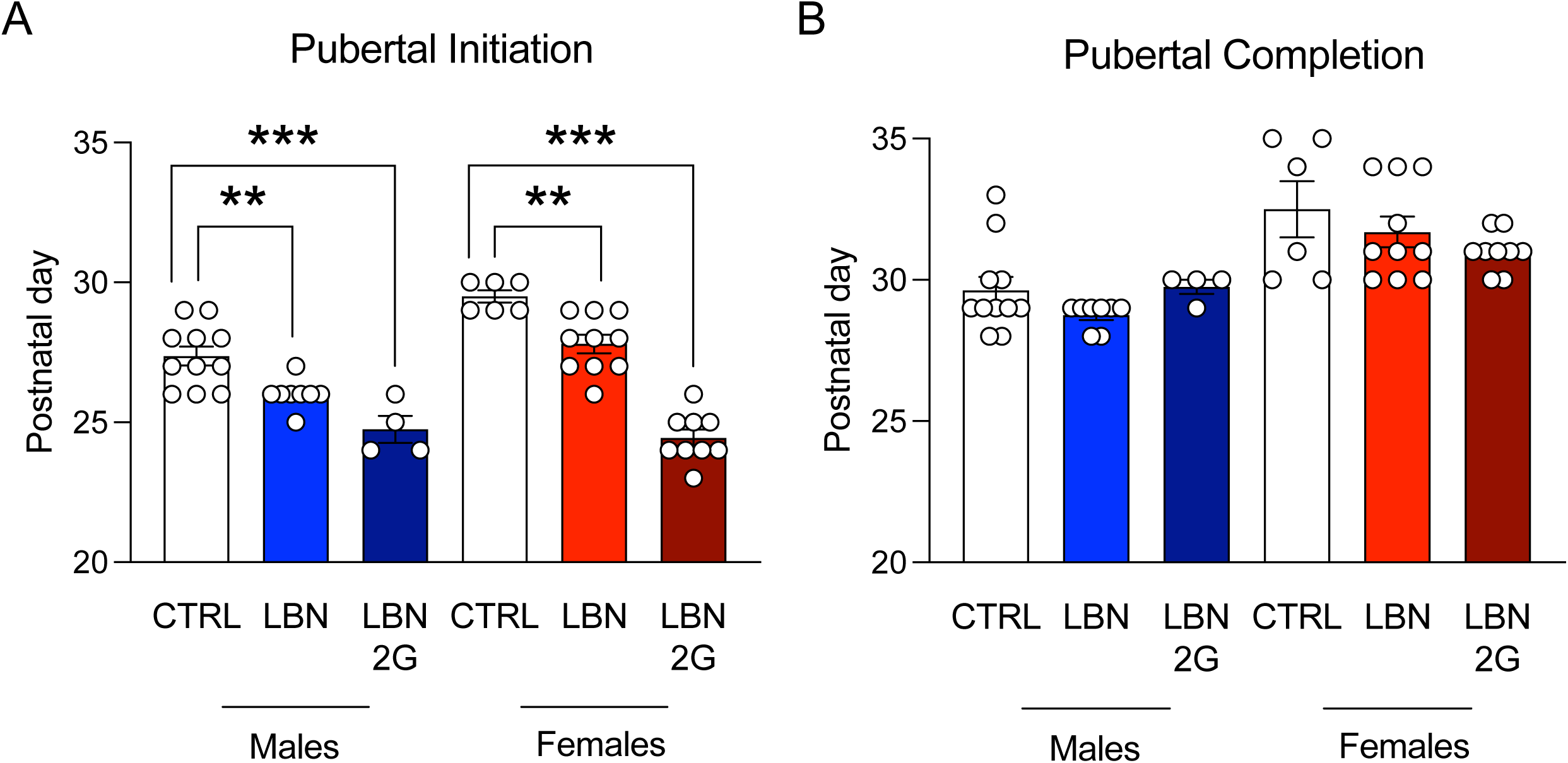
Early life stress induces early pubertal initiation in both males and females. (**A**) Pubertal initiation occurs significantly earlier in LBN and LBN 2G male and female mice as compared to CTRL mice, with a stronger effect in LBN 2G mice of both sexes. (**B**) There are no significant differences in pubertal completion in male or female mice across groups. (n = 4-11 mice per group). Error bars represent ± SEM. ** *p* < 0.01, *** *p* < 0.001. CTRL, control; LBN, Limited Bedding and Nesting; LBN 2G, LBN 2nd generation.

## DISCUSSION

Here we demonstrate that ELS, in the form of LBN, produces alterations in both the dams and their offspring that can perpetuate across generations. In the dams, our studies focused solely on behavioral endpoints, whereas in the pups we examined behavioral and physiological endpoints. LBN produced fragmented maternal behavior as well as early alterations in social communication in the offspring that did not persist into adolescence. Specifically, LBN-reared pups emitted fewer USVs as early as P3, with alterations in call length and bandwidth on P6 and P9, suggesting a higher complexity of calls as the LBN progressed and the mice matured. Since the numbers of USVs in pups are generally highest soon after birth and gradually decrease during development^40^, this outcome may indicate either early deficits in social communication or, alternatively, precocious reductions that reflect a more rapid pace of development. Although there were no effects on social behavior in adolescence, we identified robust physiological changes in peripheral systems, including sex-dependent alterations in the thymus, adrenal glands, and blood-borne levels of T-cell receptor excision circles (TRECs), a robust indication of thymic output that we have previously found to be a biomarker of prior traumatic stress in both mice and humans^18^. As was the case with USVs, some of the changes could reflect precocious development of peripheral systems that regulate stress responsiveness. Consistent with this possibility, LBN male and female mice exhibited earlier pubertal onset, suggesting significant alteration of developmental processes in response to ELS^55–57^. Remarkably, several of our findings were also seen in dams that had been exposed to LBN as well as in their LBN-exposed offspring, suggesting perpetuation of ELS effects across generations. These findings are broadly consistent with previous work using ELS models, including LBN, and studies in humans, while also raising new questions about whether early stress can have echoing effects across generations.

We first validated that LBN dams exhibited fragmented behavior, a hallmark characteristic of this paradigm^36^. We indeed confirmed that LBN dams exhibited higher behavioral transitions, indicative of fragmented maternal behavior. The LBN dams also exhibited more compulsive-like tail biting behavior, a behavior that was completely absent in CTRL dams and that has been suggested to indicate stereotypy and stress coping^60^. LBN 2G dams also exhibited a similar repertoire of fragmented behaviors, suggesting learned maternal behavior that contributes to the perpetuated effects on the pups. It was recently reported that scarcity-adversity in rats does not alter the offspring’s eventual maternal behavior when placed into standard housing conditions^61^—which we did not assess in the present studies—highlighting the unique influence of the environment in perpetuating learned maternal behaviors onto the next generation. It is important to note that by nature, the LBN paradigm only tests for the influence of maternal behavior on the offspring, without direct assessment of the influence of paternal behavior. However, both parents underwent LBN to generate perpetuated LBN 2G litters. The resulting phenotypes in the LBN 2G litters in the present study suggest the influence of both the dam’s learned maternal behavior and potential epigenetic modifications from the paternal line. This is the first study to our knowledge to test perpetuation of LBN effects by pairing two stressed parents, and having their offspring undergo the same ELS. Overall, more research is needed, including cross-fostering studies, to understand the unique contributions of learned vs. inherited behaviors from the dam to determine the roots of the perpetuated maternal behavioral phenotypes in the offspring.

We next evaluated the consequences of fragmented maternal behavior on offspring behaviors during early postnatal period and adolescence. There are no reports in the literature that assess pup social communication as early as P3 using the LBN model in mice. We uncovered that LBN rearing disrupted pup-dam communication in both the first- and second-generation litters. These findings are consistent with prior work showing that USVs are sensitive to environmental conditions and rough handling^14^ and are predictive of later social behavior in adolescence in rats^49^. In particular, the latter study found that properties of complex USVs at P10 were predictive of adolescent social play behavior. Importantly, we tested at multiple timepoints to enable repeated assessments—modeling the type of continuous data collection strategies being increasingly implemented in human studies^62^—as USVs can change over development^63^. Furthermore, while we found effects in early social communication, we did not uncover effects of LBN on adolescent social interaction except in LBN 2G male mice, suggesting that this deficit only emerges with perpetuated stress across generations. These findings contrast with prior LBN studies that have reported social deficits in adolescence^49,64^ and adulthood^65^. However, this may be due to the specific open-field SI assay used in the present study; indeed, differences in sociability reported have been more reliably detected using 2- or 3-chamber, peer play, or sexual motivation paradigms^3,36,49,64,66^. Thus, our findings may reflect both task sensitivity and sex-dependent effects. Notably, positive social buffering during early adolescence, given that the mice were placed back into normal cages and co-housed after rearing with no additional stressors, may have also temporarily masked behavioral alterations^57^. When introducing a brief restraint stress prior to SI, female LBN mice exhibited a SI deficit, consistent with studies showing LBN confers selective vulnerability to females^67^. Yet, female LBN 2G mice exhibited resistance to restraint stress, raising the possibility that epigenetic or endocrine mechanisms may transfer a transient resilience-like effect at this timepoint.

Although the early effects of LBN on social communication did not reliably persist into adolescence, we uncovered robust alterations in peripheral biology during adolescence. Increased thymus weights in LBN and LBN 2G male mice was an opposite result to what we initially hypothesized; we expected that ELS would cause thymic involution (shrinkage), indicating compromised thymic function and dysregulation of immune function. However, in mice, the thymus normally peaks in weight in adolescence (∼P40)^16,52,53^. The increased thymus size and function in LBN males led us to hypothesize that LBN induces accelerated maturation and pubertal onset, as thymus-derived signals regulate gonadal axis maturation, causing the thymus weight to peak earlier than CTRL counterparts. Intriguingly, while thymus size increased in LBN 2G male mice, there was not a correspondingly expected increase in TREC levels; similarly for female LBN 2G mice, though there were no changes in thymus size, TREC levels increased. We have previously reported TRECs as a marker of prior traumatic stress in mice and humans, and that they are related to epigenetic regulation of the innate immune response^18^, providing a potential avenue of further exploration for these disparate and sex-dependent effects. When we introduced a restraint stressor, the thymus involuted in LBN 2G females, reflecting the thymus’s known sensitivity to restraint stress^68^. Similarly, while adrenal size was not significantly impacted across groups, we found blunting of CORT and exacerbation of IL-6 in blood of LBN 2G male and female mice. Previous work has shown that stress effects on adrenals and thymus can be uncoupled^18^, raising the possibility that multiple mechanisms are involved in these changes.

Considered together, our data highlight that early ELS-induced communication deficits may not persist behaviorally but do leave a lasting biological signature on immune and endocrine systems that may suggest accelerated biological aging. Specifically, our discovery that LBN induces thymic alterations alongside earlier pubertal initiation aligns with theories of accelerated biological aging after ELS^57,69^. Supporting this, ELS has been linked to shortened telomere length^70^ and epigenetic aging signatures, with especially strong effects in females during puberty, a window of heightened disease risk. Several rodent studies have also reported accelerated maturation of sensory and motor development^71^ as well as the hippocampus^55^ and amygdala^66,67^. One study similarly reported earlier pubertal initiation but not completion in female mice after maternal separation (MS)^49^, while another found female-specific maturation but a delay in males after LBN in rats^72^. We hypothesize that this accelerated maturation may offer short-term survival benefits in unpredictable environments by promoting earlier independence, but at the cost of long-term vulnerability to psychiatric and physical disease later in life. However, the Bath lab has found delayed sexual maturation in female mice using the LBN (conducted on P4-P11)^58^ and has also reported variable effects of ELS on maturation depending on the timepoint and biological endpoint measured (e.g., somatic growth, eye opening)^59^. These data highlight the importance of studying the longitudinal effects of ELS on maturational milestones and determining whether they are specific biomarkers of future stress susceptibility. Future work will assess stress sensitivity across the lifespan to determine vulnerability later in life as well as the rate of biological aging by assessing the biological clock.

This study also demonstrates that LBN induces early-life communication deficits and lasting biological alterations that extend intergenerationally. Specifically, our LBN 2G data also reflect human intergenerational trauma studies demonstrating blunted corticosterone (CORT) responses^21,73–75^. Ongoing and future studies will probe epigenetic mechanisms underlying LBN 2G resilience, adaptation, and accelerated aging and incorporate adoption designs to parse inherited versus maternal behavioral transmission. Treatments to modify peripheral adaptations in early age to prevent any intergenerational stress perpetuation effects will also be pursued. Importantly, the findings in the present study are consistent with observations described in the human literature: earlier pubertal onset, faster biological aging, and long-term susceptibility to illness across the lifespan leading to reduced healthspan, or the number of years lived free of chronic illness^76^. Overall, the present study elucidates potential biomarkers to predict outcomes, enable proactive intervention, and reduce the potentially harmful effects of stress on healthspan.

## ACKNOWLEDGMENTS

We thank members of the Carlezon and Ressler laboratories for helpful feedback on the experimental design and manuscript. We thank Dr. Heather Brenhouse for sharing and demonstrating her laboratory’s pubertal tracking protocol.

## AUTHOR CONTRIBUTIONS

JCM: Conceptualization, Methodology, Investigation, Data Analyses, Writing (Original Draft and Editing); JLK: Methodology, Data Analyses, Writing (Original Draft); NSC: Methodology, Data Analyses; CWS: Data Analyses; TO (Temi): Data Analyses; TO (Tolu): Data Analyses; TZB: Original Methodology, Writing (Review and Editing); OOF: Conceptualization, Methodology, Investigation, Writing (Review and Editing); WAC: Conceptualization, Funding Acquisition, Writing (Review and Editing).

## DATA AVAILABILITY

### Lead contact

Further information and requests for resources should be directed to and will be fulfilled by Dr. William (Bill) A. Carlezon, Jr.

### Data and materials availability

All reported data collected in this study and code used for analysis are available upon request from the corresponding authors. Any additional information required to reanalyze the data reported in this paper is available from the lead contact upon request. This study did not generate new unique reagents.

