## Supplemental Information for "Early life stress induces social behavioral deficits and peripheral biomarker alterations in adolescence that perpetuate intergenerationally"

Corresponding author emails:

### SUPPLEMENTAL METHODS

*Ultrasonic vocalization (USV) recording and analysis*

USVs were recorded for each pup on P3, P6, and P9 to capture alterations in social behavior throughout rearing conditions. The dam was separated from her litter for 10 min before recording. Each pup was recorded separately in a Styrofoam sound chamber. Because decreased body temperature can induce USVs, the cage was placed above a heating pad at 37°C to maintain body temperature during experimentation. An ultrasonic microphone (Avisoft Bioacoustics, model CM16/CMPA) positioned 10 cm above the Styrofoam sound chamber was used to record the pups for a 3-min recording period. Following the recording, the pup was returned to the home cage.

Audio files were uploaded and analyzed using DeepSqueak [1], a Matlab program for USV analysis. USVs were visualized by converting each file to spectrograms using Fast Fourier Transformation. Each call detected by DeepSqueak was manually confirmed or rejected by a trained experimenter blinded to experimental conditions. The total number of calls, duration, maximum frequency, minimum frequency, and peak frequency of each call were determined. Bandwidth was calculated by the difference between maximum and minimum frequency. For each pup, the duration and bandwidth were averaged across all vocalizations.

*Social interaction (SI)*

An SI paradigm was conducted at P30 to determine alterations in social behavior at a critical adolescent timepoint [2–5]. Behavior occurred during the animal's light cycle in the mornings. Mice were placed into a large open field containing 1 upside-down wire mesh pencil cups. During the first trial, the pencil cup served as a novel object or empty container. Mice then underwent an intertrial period of 1 min. During the second trial, the pencil cup contained a novel adult BALB/cJ non-aggressive male or ovariectomized female mouse, to match the sex of the experimenter mouse. For each trial, mice were placed in the middle of the open field arena and allowed to explore for 2.5 min. Sessions were videotaped and were later analyzed using behavioral tracking software (Noldus EthoVision, Leesberg, VA). Two hours after the start of SI, mice were euthanized; blood and tissues were collected for qPCR as described [6] and adrenal and thymus glands were dissected, cleaned for excess tissue or fat, and weighed. Organ weights were calculated as milligrams per kilogram (mg/kg) of body weight.

*Restraint stress*

A separate group of mice underwent a 10 min restraint stress 2 h before the start of SI. In brief, mice were placed in decapicones specifically designed for mouse restraint (Braintree Scientific Inc., Braintree, MA) for 10 min, then placed back into their home cages. Two hours later, mice underwent the SI paradigm and were then euthanized as described above.

*Quantitative polymerase chain reaction (qPCR)*

For quantitative analyses, real-time (RT)-PCR was performed as previously described [6–8] using an Applied Biosystems ViiA7 Real-Time PCR System. Primer sequences are listed in **Supplemental Table 2**. Specific primers were used for the TREC signal joint in C57BL/6 J mice, and to compensate for input, the constant segment of C57BL/6J T-cell receptor alpha (TCRA) gene was measured as reference [7]. Changes (delta, [d]) in cycle threshold (dCT) were calculated as CtTREC-CtTCRA, and only calculated for samples whose CtTREC was <35 with replicate standard deviation <0.5 Ct. For clarity in depicting the data, values are expressed as -dCT, with larger integer values indicating lower TREC levels.

### SUPPLEMENTAL FIGURE LEGENDS

**Supplemental Figure 1. Maternal behavior is reliably fragmented throughout the rearing period using the LBN model and transmits intergenerationally.** (**A-B**) LBN and LBN 2G dams showed a significant increase in both duration and counts of low nursing behaviors. (**C-D)** There was no difference in time the dam spent off nest, but an increase in counts for the LBN and LBN 2G groups. (**E**) Total behavioral event counts of all behaviors were significantly higher in the LBN and LBN 2G groups. (**F**) The average licking and grooming pups (LG) duration was reduced in LBN and LBN 2G groups. (**G**) Despite the decreased average duration of LG in the LBN and LBN 2G dams, there was no difference in total LG duration across groups. (**H**) However, there is an increase counts of LG in LBN and LBN 2G dams. (**I-J**) LBN 2G dams displayed decreased side nursing duration, while there were no differences in side nursing counts across groups. (**K-L**) Dams in LBN conditions exhibited greater tail biting behavior duration and counts. (**M-N**) No differences were observed in eating behavior duration, though LBN dams demonstrated a trend towards an increase in counts of eating. (**O-P**) There were no differences were observed across groups in duration or counts of active nursing. (**Q-R**) No significant differences were observed in duration or counts of carrying pups. (**S-T**) LBN 2G dams exhibited a trend towards reduced duration and counts of nestbuilding compared to both CTRL and the LBN dams. (**U-V**) LBN and LBN 2G dams exhibited higher counts of moving on the nest. (**W-X**) LBN and LBN 2G dams exhibited lower self-grooming duration with no difference in counts. (**Y-Z**) When moving on nest and off nest durations and counts were summed, no differences in duration were observed across groups. However, both LBN and LBN 2G dams exhibited increased counts. (**AA-BB**) No differences were observed across groups in duration of pup-directed behaviors, but the LBN and LBN 2G dams showed a significant increase in counts. (**CC-DD**) There were no noticeable differences in duration or counts of self-directed behavior across groups. (n = 6 dams per group). Error bars represent ± SEM. * $p < 0.05$, ** $p < 0.01$, *** $p < 0.001$. CTRL, control group; LBN, Limited Bedding and Nesting group; LBN 2G, LBN 2nd generation; M, Moving on Nest; O, Off the Nest, LG, Licking and Grooming Pups; sec, seconds.

**Supplemental Figure 2. Summary of maternal behavior during the inactive period.** (**A-B**) No significant differences were observed in duration or counts of low nursing during the inactive phase. (**C-D**) There were no recorded differences in the duration or counts of the dam off the nest during the inactive phase. (**E**) There were significantly higher total event counts during the inactive phase in both LBN and LBN 2G dams. (**F**) LBN dams exhibited decreased average licking and grooming (LG) duration. (**G-H**) There were no differences in LG duration or counts across all groups. (**I-J**) LBN 2G dams exhibited decreased duration of side nursing. (**K-L**) In the inactive phase, the tail biting behavior was evident in both LBN and LBN 2G groups. (**M-N**) No significant differences appeared in eating duration or counts. (**O-P**) No significant differences were recorded in active nursing duration or counts. (**Q-R**) LBN 2G dams exhibited higher duration of carrying pups. (**S-T**) No significant differences appeared in nestbuilding duration or counts, but the LBN 2G group demonstrated no instances of nestbuilding. (**U-V**) LBN and LBN 2G dams exhibited higher counts of moving on the nest. (**W-X**) No significant differences appeared in self-grooming duration or counts. (**Y-**

**Z**) LBN and LBN 2G dams exhibited significantly higher counts of moving on nest and off the nest. (**AA-BB**) No significant differences appeared in pup-directed duration or counts. (**CC-DD**) No significant differences were recorded in self-directed duration or counts. (**EE-FF**) LBN and LBN 2G dams exhibited pups off the nest during the entire inactive phase. (**GG-II**) Representative images of homecage behavior demonstrating: (**GG**) a CTRL dam laying on her nest with her pups; (**HH**) an LBN dam eating facing away from the nest; and (**II**) an LBN 2G dam tail biting. (n = 6 dams per group). Error bars represent ± SEM. * $p < 0.05$, ** $p < 0.01$, **** $p < 0.0001$. CTRL, control group; LBN, Limited Bedding and Nesting group; LBN 2G, LBN 2nd generation; M, Moving on Nest; O, Off the Nest, AN, Active Nursing; C, Carrying Pups; E, Eating; LG, Licking and Grooming Pups; LN, Low Nursing; NB, Nestbuilding; SG, Self-Grooming; SN, Side Nursing; TB, Tailbiting; sec, seconds.

**Supplemental Figure 3. Summary of maternal behavior during the active period.** (**A-B**) LBN 2G dams exhibited an increase in duration and counts of low nursing. (**C-D**) For dam off the nest, no significant differences were observed in duration, with the LBN 2G group exhibiting a trend towards an increase in counts. (**E**) For total behavioral event counts for the active phase on P6, both the LBN and LBN 2G groups exhibited increased counts compared to the CTRL group. (**F**) LBN 2G dams had a significantly decreased average duration of LG than the CTRL dams. (**G-H**) No significant differences were observed across licking and grooming pups duration or counts, though LBN 2G dams trended towards an increase in counts. (**I-J**) There were no significant differences in duration of side nursing, but a trending increase in counts in LBN dams.

(**K-L**) Tail biting duration and counts were significantly increased in LBN and LBN 2G dams. (**M-N**) There were no differences in duration or counts of eating across groups. (**O-P**) No significant differences in duration or counts of active nursing were observed across groups. (**Q-R**) There were no differences observed across groups in carrying pups. (**S-T**) LBN dams demonstrated a trend towards an increase in nest building. (**U-V**) For moving on nest, the LBN 2G group showed an increase in counts. (**W-X**) Self grooming behavior duration and counts decreased in LBN and LBN 2G dams. (**Y-Z**) There were differences across groups in duration of moving on and off the nest, though LBN 2G group exhibited a trend towards increase in counts of these behaviors. (**AA-BB**) No differences were observed in pup-directed behavior duration (nursing, carrying pups, nestbuilding, and licking and grooming pups), but both LBN and LBN 2G mice exhibited a significant increase in counts. (**CC-DD**) No significant differences were observed for self-directed behavior, though the LBN 2G group exhibited a trend towards a decrease in duration. (**EE-FF**) LBN and LBN 2G pups were off the nest during the entire active phase, while CTRL pups stayed on the nest. (**GG-II**) Representative images of homecage behavior demonstrating: (**GG**) a CTRL dam nursing her pups in a cage with normal bedding and nesting materials; (**HH**) an LBN dam facing away from her nest; and (**II**) an LBN 2G dam with pups off the nest. (n = 6 dams per group). Error bars represent ± SEM. * $p < 0.05$, ** $p < 0.01$, *** $p < 0.001$. CTRL, control group; LBN, Limited Bedding and Nesting group; LBN 2G, LBN 2nd generation; M, Moving on Nest; O, Off the Nest, AN, Active Nursing; C, Carrying Pups; E, Eating; LG, Licking and Grooming Pups; LN, Low Nursing; NB, Nestbuilding; SG, Self-Grooming; SN, Side Nursing; TB, Tailbiting; sec, seconds.

**Supplemental Figure 4. The number of pups per litter does not differ across maternal conditions.** The number of pups per litter does not differ across CTRL, LBN, and LBN 2G mice included in the present study. CTRL, control; LBN, limited bedding and nesting; LBN 2G, LBN 2$^{nd}$ generation.

**Supplemental Figure 5. Representative spectrograms of ultrasonic vocalizations in pups.** (**A-C**) Example spectrograms taken from DeepSqueak representing CTRL, LBN, and LBN 2G calls during P9 over a 1s time period. CTRL, control; LBN, limited bedding and nesting; LBN 2G, LBN 2$^{nd}$ generation; P, postnatal day; USV, ultrasonic vocalization; s, second.

**Supplemental Figure 6. Extended data of social interaction in adolescence from Figure 3.** (**A**) There is no difference in time spent with the empty cup across groups in males. (**B**) There is no difference in time spent with the empty cup across groups in females. (**C**) There is no difference in overall time spent with the mouse across groups in males. (**D**) There is no difference in overall time spent with the mouse across groups in females. (**E**) There is no difference in total distance traveled across groups in males. (**F**) There is no difference in total distance traveled across groups in females. (**G**) There is no difference in average velocity across groups in males. (**H**) There is no difference in average velocity across groups in females. (n = 4-20 mice per group). Error bars represent ± SEM. * $p < 0.05$. CTRL, control group; LBN, Limited Bedding and Nesting group; LBN 2G, LBN 2nd generation; No., number; sec, seconds; cm, centimeters.

**Supplemental Figure 7. The number of ultrasonic vocalizations is differentially altered depending on postnatal day, sex, and rearing environment.** (**A**) LBN mice call less than CTRL mice as early as P3, with LBN 2G mice trending towards lower calls, replicating data from **Figure 2**. (**B**) USV number on P6 trends lower in LBN and LBN 2G male mice and is significantly lower in LBN female mice as compared to CTRL mice. (**C**) USV number on P9 is lower in LBN males and trends lower in LBN 2G males as compared to CTRL males, whereas LBN 2G females call more on P9 as compared to CTRL females. (**D**) Total USV number across all postnatal days tested is significantly lower in LBN male and female mice as compared to their CTRL counterparts but is not altered in LBN 2G mice. (n = 7-22 mice per group). Error bars represent ± SEM. * $p < 0.05$; ** $p < 0.01$; *** $p < 0.001$. CTRL, control; LBN, limited bedding and nesting; LBN 2G, LBN 2$^{nd}$ generation; USV, ultrasonic vocalization; P, postnatal day.

**Supplemental Figure 8. Extended data of social interaction in adolescence from Figure 4.** (**A**) There is no difference in time spent with the empty cup across groups in males. (**B**) There is no difference in time spent with the empty cup across groups in females. (**C**) There is no difference in overall time spent with the mouse across groups in males. (**D**) There is no difference in overall time spent with the mouse across groups in females. (**E**) There is no difference in male bouts to the mouse across groups in males. (**F**) There is no difference in bouts to the mouse across groups in females. (**G**) There is no difference in total distance traveled across groups in males. (**H**) There is no difference in total distance traveled across groups in females. (I) There is no difference in average velocity across groups in males. (**J**) There is no difference in average velocity across groups in females. (n = 6-19 mice per group). Error bars represent ± SEM. CTRL, control group; LBN, Limited Bedding and Nesting group; LBN 2G, LBN 2nd generation; No., number; sec, seconds; cm, centimeters.

**Supplemental Figure 9. Non-normalized thymus and adrenal weights across groups.** (**A**) Thymus size not normalized over mouse body weight does not differ across groups in males. (**B**) Adrenal size not normalized over mouse body weight does not differ across groups in males. (**C**) Thymus weights of LBN 2G mice not normalized by mouse body weight is lower than those of CTRL female mice. (**D**) Adrenal size not normalized over mouse body weight does not differ across groups in females. (n = 10-28 mice per group). Error bars represent ± SEM. ** $p < 0.01$. CTRL, control; LBN, limited bedding and nesting; LBN 2G, LBN 2$^{nd}$ generation; mg, milligram.

**Supplemental Figure 10. CORT and IL-6 are altered across groups in a generational- and sex-dependent manner.** (**A**) LBN male mice exhibit an increase in plasma CORT expression as compared to CTRL male mice, which is blunted in LBN 2G mice. Female mice do not exhibit alterations in CORT across groups. (**B**) Plasma IL-6 expression trends higher in both LBN and LBN 2G males as compared to CTRL males. IL-6 also trends higher in LBN females as compared to CTRL females and is significantly higher in LBN 2G females. CTRL, control; LBN, limited bedding and nesting; LBN 2G, LBN 2$^{nd}$ generation; pg/mL, picogram per milliliter; CORT, corticosterone; IL-6, interleukin-6.

**Supplemental Figure 11. Extended data of social interaction in adolescence from Figure 6.** (**A**) There is no difference in time spent with the empty cup across groups in males. (**B**) There is no difference in time spent with the empty cup across groups in females. (**C**) There is no difference in overall time spent with the mouse across groups in males. (**D**) LBN females spend significantly less time with the mouse than CTRL females. (**E**) LBN 2G males trended towards significantly lower bouts to the mouse than CTRL males. (**F**) There is no difference in bouts to the mouse across groups in females. (**G**) There is no difference in total distance traveled across groups in males. (**H**) There is no difference in total distance traveled across groups in females. (**I**) There is no difference in average velocity across groups in males. (**J**) There is no difference in average velocity across groups in females. (n = 3-22 mice per group). Error bars represent ± SEM. * $p < 0.05$. CTRL, control group; LBN, Limited Bedding and Nesting group; LBN 2G, LBN 2nd generation; No., number; sec, seconds; cm, centimeters.

**Supplemental Figure 12. Non-normalized thymus and adrenal weights across restraint groups.** (**A**) LBN 2G male mice have greater thymus size when not normalized over mouse body weight. (**B**) Adrenal size not normalized over mouse body weight does not differ across groups in males. (**C**) Thymus weights of LBN 2G mice not normalized by mouse body weight is lower than those of CTRL female mice. (**D**) Adrenal size not normalized over mouse weight trends lower in LBN female mice and is significantly lower in LBN 2G female mice. (n = 3-21 mice per group). Error bars represent ± SEM. ** $p < 0.01$; *** $p < 0.001$. CTRL, control; LBN, limited bedding and nesting; LBN 2G, LBN 2nd generation; mg, milligram.

**Supplemental Figure 13. Example images of thymus and adrenals in adolescence after early life stress and across generations in males and females.** (**A-C**) Representative thymus photos from male CTRL, LBN, and LBN 2G mice across experiments. (**D-F**) Representative adrenal photos from female CTRL, LBN, and LBN 2G mice that underwent restraint stress. CTRL, control; LBN, limited bedding and nesting; LBN 2G, LBN 2nd generation.

**Supplemental Table 1. Statistical analyses.**

**Supplemental Table 2. Primer sequences utilized for TREC RT-qPCR.**

**Supplemental Video 1 – Example LBN dam exhibiting tail biting behavior.**

**Supplemental Video 2 – Example control dam behavior at postnatal day 6.**

**Supplemental Video 3 – Example LBN dam tail chasing behavior at postnatal day 6.**

**Supplemental Video 4 – Example LBN 2G dam behavior at postnatal day 6.**
