## Supplemental Figure 5 for "Early life stress induces social behavioral deficits and peripheral biomarker alterations in adolescence that perpetuate intergenerationally"

A) CTRL Vocalization Spectrogram @ P9

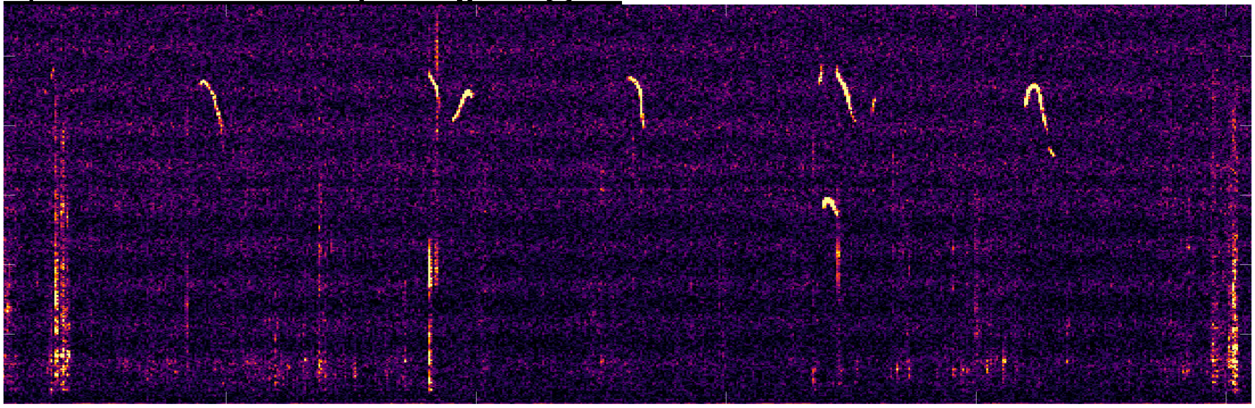

(over 1s time period)

B) LBN Vocalization Spectrogram @ P9

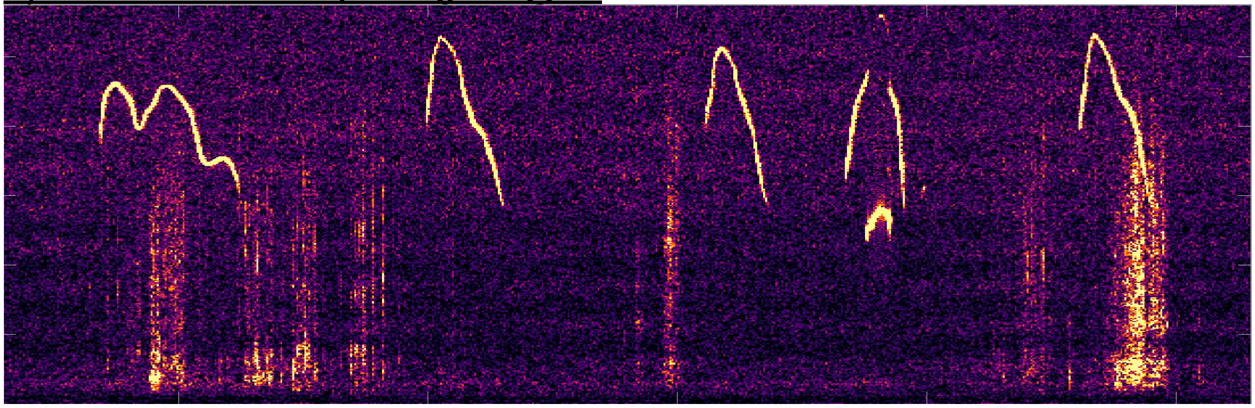

(over 1s time period)

C) LBN 2nd Generation Vocalization Spectrogram @ P9

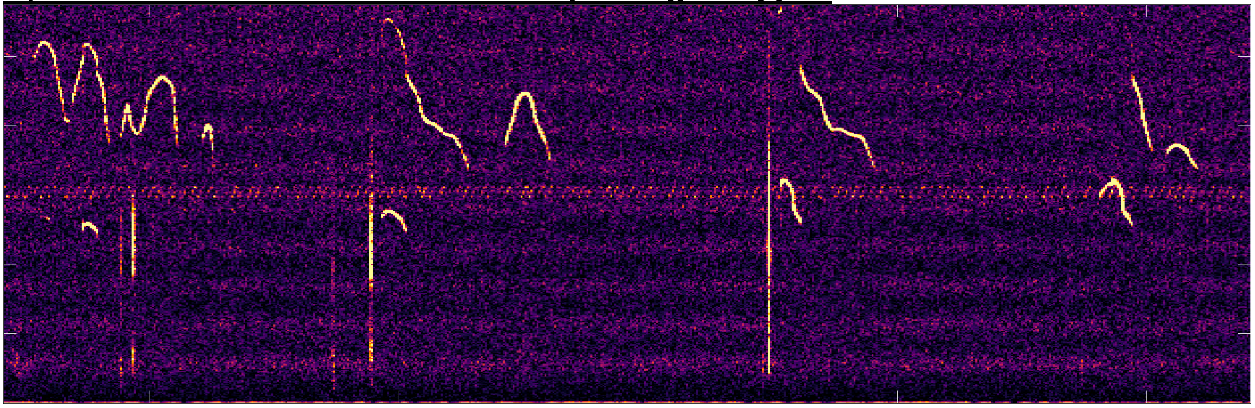

(over 1s time period)
